# NAC1 directs CEP1-CEP3 peptidase expression and modulates root hair growth in Arabidopsis

**DOI:** 10.1101/2023.03.15.532802

**Authors:** Diana R. Rodríguez-García, Yossmayer del Carmen Rondón Guerrero, Lucía Ferrero, Andrés Hugo Rossi, Esteban A. Miglietta, Ariel A. Aptekmann, Eliana Marzol, Javier Martínez Pacheco, Mariana Carignani, Victoria Berdion Gabarain, Leonel E. Lopez, Gabriela Díaz Dominguez, Cecilia Borassi, José Juan Sánchez-Serrano, Lin Xu, Alejandro D. Nadra, Enrique Rojo, Federico Ariel, José M. Estevez

**Affiliations:** Fundación Instituto Leloir and IIBBA-CONICET. Av. Patricias Argentinas 435, Buenos Aires C1405BWE, Argentina; Instituto de Agrobiotecnología del Litoral, CONICET, Universidad Nacional del Litoral, Colectora Ruta Nacional 168 km 0, 3000, Santa Fe, Argentina; Departamento de Fisiología, Biología Molecular y Celular, Instituto de Biociencias, Biotecnología y Biología Traslacional (iB3). Facultad de Ciencias Exactas y Naturales, Universidad de Buenos Aires, Ciudad Universitaria, Buenos Aires C1428EGA, Argentina; Departamento de Química Biológica, Facultad de Ciencias Exactas y Naturales, Universidad de Buenos Aires (IQUIBICEN-CONICET), Ciudad Universitaria, Buenos Aires C1428EGA, Argentina; National Laboratory of Plant Molecular Genetics, CAS Center for Excellence in Molecular Plant Sciences, Institute of Plant Physiology and Ecology, Shanghai Institutes for Biological Sciences, Chinese Academy of Sciences, Shanghai 200032, China; Centro Nacional de Biotecnología, Consejo Superior de Investigaciones Científicas, Cantoblanco, E-28049 Madrid, Spain; Centro de Biotecnología Vegetal, Facultad de Ciencias de la Vida, Universidad Andrés Bello and Millennium Institute for Integrative Biology (iBio), Santiago, Chile; ANID – Millennium Institute for Integrative Biology (iBio), 7500000 Santiago, Chile; ANID –Millennium Nucleus for the Development of Super Adaptable Plants (MN-SAP), 8331150 Santiago, Chile

**Author notes:** Correspondence should be addressed. (J.M.E). Author for Correspondence: José M. Estevez Fundación Instituto Leloir, Av. Patricias Argentinas 435, Buenos Aires C1405BWE, Argentina. TE: 54-115238-7500 EXT. 3206 Centro de Biotecnología Vegetal, Facultad de Ciencias de la Vida, Universidad Andrés Bello and Millennium Institute for Integrative Biology (iBio), Santiago CP 8370146, Chile.

## Abstract

Plant genomes encode a unique group of papain-type Cysteine EndoPeptidases (CysEPs) containing a KDEL endoplasmic reticulum (ER) retention signal (KDEL-CysEPs or CEPs). CEPs process the cell-wall scaffolding EXTENSIN proteins (EXTs), which regulate *de novo* cell wall formation and cell expansion. Since CEPs are able to cleave EXTs and EXT-related proteins, acting as cell wall-weakening agents, they may play a role in cell elongation. *Arabidopsis thaliana* genome encodes three CEPs (AtCPE1-AtCEP3). Here we report that the three Arabidopsis CEPs, *AtCEP1-AtCEP3,* are highly expressed in root-hair cell files. Single mutants have no evident abnormal root-hair phenotype, but *atcep1-3 atcep3-2* and *atcep1-3 atcep2-2* double mutants have longer root hairs (RHs) than wild type (Wt) plants, suggesting that expression of *AtCEPs* in root trichoblasts restrains polar elongation of the RH. We provide evidence that the transcription factor *NAC1* activates AtCEPs expression in roots to limit RH growth. Chromatin immunoprecipitation indicates that NAC1 binds the promoter of *AtCEP1, AtCEP2,* and to a lower extent to *AtCEP3* and may directly regulate their expression. Indeed, inducible *NAC1* overexpression increases *AtCEP1 and AtCEP2* transcript levels in roots and leads to reduced RH growth while the loss of function *nac1-2* mutation reduces *AtCEP1-AtCEP3* gene expression and enhances RH growth. Likewise, expression of a dominant chimeric NAC1-SRDX repressor construct leads to increased RH length. Finally, we show that RH cell walls in the *atcep1-1 atcep3-2* double mutant have reduced levels of EXT deposition, suggesting that the defects in RH elongation are linked to alterations in EXT processing and accumulation. Taken together, our results support the involvement of AtCEPs in controlling RH polar growth through EXT-processing and insolubilization at the cell wall.

## Introduction

In plants there is a unique group of papain-type Cysteine EndoPeptidases (CysEPs) containing a KDEL endoplasmic reticulum (ER) retention signal (KDEL-CysEPs or CEPs) for which no homologous genes have been identified in mammals or yeast (Gietl et al. 2000). CEPs are synthesized as pre-pro-enzyme and co-translationally translocated into the ER, where the pre-sequence is removed and the pro-enzyme is finally released from the ricinosomes (ER-associated structures) upon vacuolar collapse and acidification of the cytosol, triggering the maturation of the enzyme (Schmid et al. 1999, 2001; Beers et al. 2000; Zhang et al. 2014). AtCEP1 (At5g50260), AtCEP2 (At3g48340), and AtCEP3 (At3g48350) are three *Arabidopsis thaliana* AtCEPs with broad expression patterns in vegetative and reproductive tissues along the plant, including root tissues (Helm et al. 2008; Hierl et al. 2014; Zhou et al. 2016). Several CEPs have been identified in cells or tissues associated with programmed cell death (PCD), where they play crucial roles in intracellular protein degradation (Tanaka 1991; Becker et al. 1997; He and Kermode 2003; Zhang et al. 2014; Olvera-Carrillo Y et al 2015; Chen et al. 2016). AtCEP1 was found to be a central mediator of tapetal PCD, allowing tapetal cell degeneration and functional pollen formation (Zhang et al. 2014). In addition, AtCEP1 and AtCEP2 in Arabidopsis were found to be functional in the *de novo* emergence of adventitious root tips associated with EXT-degradation and regulated by the NAC1 transcription factor (Chen et al. 2016). More recently, it was reported that *AtCEP1* and *AtCEP2* are expressed in root epidermal cells that separate to allow LR emergence, and that loss of function of *AtCEP1* or *AtCEP2* causes delayed emergence of LR primordia, suggesting that these KDEL-CysEPs might be involved in cell-wall remodelling for cell separation during development (Howing et al. 2018).

In addition to the papain-type preference for neutral amino acids with large aliphatic and non-polar (Leu, Val, Met) or aromatic (Phe, Tyr, Trp) side chains in the P2 position, Ricinus CEP (RcCysEP) exhibits an unusually broad substrate specificity. This broad substrate specificity is a result of the active site cleft of KDEL-CysEPs, which accepts a wide variety of amino acids, including proline and the glycosylated hydroxyproline of hydroxyproline-rich glycoproteins (HRGP) of the cell wall (Than et al. 2004). The amino acid residues that are essential for this generally more open structure of the active site cleft, as well as those that define the catalytic pocket, are highly conserved among known KDEL-CysEPs (Hierl et al. 2014). It can recognize the Ser-(Hyp)_3-5_ repeats, *O*-glycosylated Hyp, and prolines at one-two amino acids relative to the cleavage site (Than et al., 2004; Helm et al. 2008; Hierl et al. 2014). These Ser-(Hyp)_3-5_ repeats with *O*-glycosylated modifications are frequently observed in structural *O*-glycoproteins Extensins (EXTs) and possibly in a large number of uncharacterized apoplastic EXT-related proteins (e.g. PERK, Formins, AGP-EXTs hybrids) (Borassi et al. 2016). 59 encoded EXTs in *Arabidopsis thaliana* contain a Tyr-crosslinking motif close to an *O*-glycosylated Ser-(Hyp)_3-5_ (Showalter et al., 2010; Marzol et al. 2018). EXT Tyr-mediated crosslinking is catalyzed by apoplastic peroxidases (Schnabelrauch et al., 1996; Jackson et al., 2001) and allows them to form glycoprotein networks in the cell wall, which influences de novo plant cell wall formation (Cannon et al., 2008) and in polar cell expansion processes (Velasquez et al. 2011; 2015).

Recently, it was shown that AtEXT3/RSH is not essential for early embryogenesis or plant viability suggesting that its function is likely to be redundant with other related EXT proteins (Doll et al. 2022). Since CEPs can cleave *O*-glycosylated EXTs (Hierl et al., 2014), thus acting as cell wall-weakening agents, this lends credence to the idea that CEPs may play a pivotal role in cell elongation. Prior to this study, we determined that at least six EXTs (EXT6–7, 12–14, 18) co-regulated at the transcriptional level play a crucial role in the polar-cell expansion process, specifically in RHs in Arabidopsis (Velasquez et al. 2011; Velasquez et al 2015; Marzol et al. 2018) and in Tomato (Bucher et al. 1997; 2002). In this process, Leucine Rich Extensins 1 and 2 (LRX1 and LRX2) were also found to be essential (Baumberger 2001, 2003; Ringli 2010). Pollen EXTs (PEXs) and LRXs were also recently linked to polar growth regulation in pollen tubes (Sede et al. 2018; Ndinyanka Fabrice et al., 2017; Wang et al., 2018), highlighting a conserved role of these EXT and EXT-related proteins in polar growth. On the basis of these previous findings, we hypothesized that Arabidopsis AtCEPs may play a role in polar-growth regulation linked to the processing of *O*-glycosylated EXT and EXT-related proteins processing and possibly other substrates during their maturation along the secretory pathway. Here, we provide evidence that, indeed, AtCEPs negatively regulate levels of EXT secretion/insolubilization at the cell-wall of root hairs (RHs) and restrain their growth.

## Results and Discussion

Expression of Arabidopsis AtCEPs has been previously analysed using functional reporter constructs that include the coding sequences of AtCEP1 and AtCEP2 under the control of their respective regulatory regions of 2.0 Kb fragment upstream of the start codon, with HA and fluorescent protein tags, either GFP or mCherry (AtCEP1pro:PRE-PRO-3xHA-EGFP-AtCEP1-KDEL and AtCEP2pro:PRE-PRO-3xHA-mCherry-AtCEP2-KDEL) inserted between the pro-peptide and the mature enzyme sequence (Howing et al. 2014 and Hierl et al. 2014). Intriguingly, it was reported that AtCEP2pro:PRE-PRO-3xHA-mCherry-AtCEP2-KDEL is expressed specifically in non-protruding cell files of the hypocotyl, the cell files where stomata are formed, but not in protruding cell files (Hierl et al., 2014). Interestingly, the alternating pattern of non-protruding and protruding cell files in the hypocotyl is controlled by common mechanisms to the alternating pattern of root-hair cell (root trichoblast) and non-root-hair cell (root atrichoblast) files in the root. Hypocotyl non-protruding cells and root trichoblasts are always positioned over an anticlinal cell wall of the underlying cortex and lack expression of cell fate regulators of GL2 and WER and the enhancer-trap marker J2301, whereas hypocotyl protruding cells and root atrichoblast are positioned over a periclinal cortex cell wall and express the J2301 marker and GL2 and WER, which block development of stomata or RHs in these cell files (Berger et al.,1998, Grierson et al., 2014). Importantly, we found that AtCEP2pro::PRE-PRO-3xHA-mCherry-AtCEP2 is expressed in root epidermis exclusively in the trichoblasts (**Figure 1A-B**; **Figure S1**), further supporting that root-hair cell files share specification mechanisms and characteristics with non-protruding cell files of the hypocotyl (Berger et al.,1998, Grierson et al., 2014). AtCEP1pro:PRE-PRO-3xHA-EGFP-AtCEP1 was also specifically expressed in root trichoblasts (**Figure 1A-B**; **Figure S1**). In these cells, the mCherry-AtCEP2 fusion protein was found in mobile punctate structures, consistent with it being localized in the endomembrane system.

**Figure 1.**
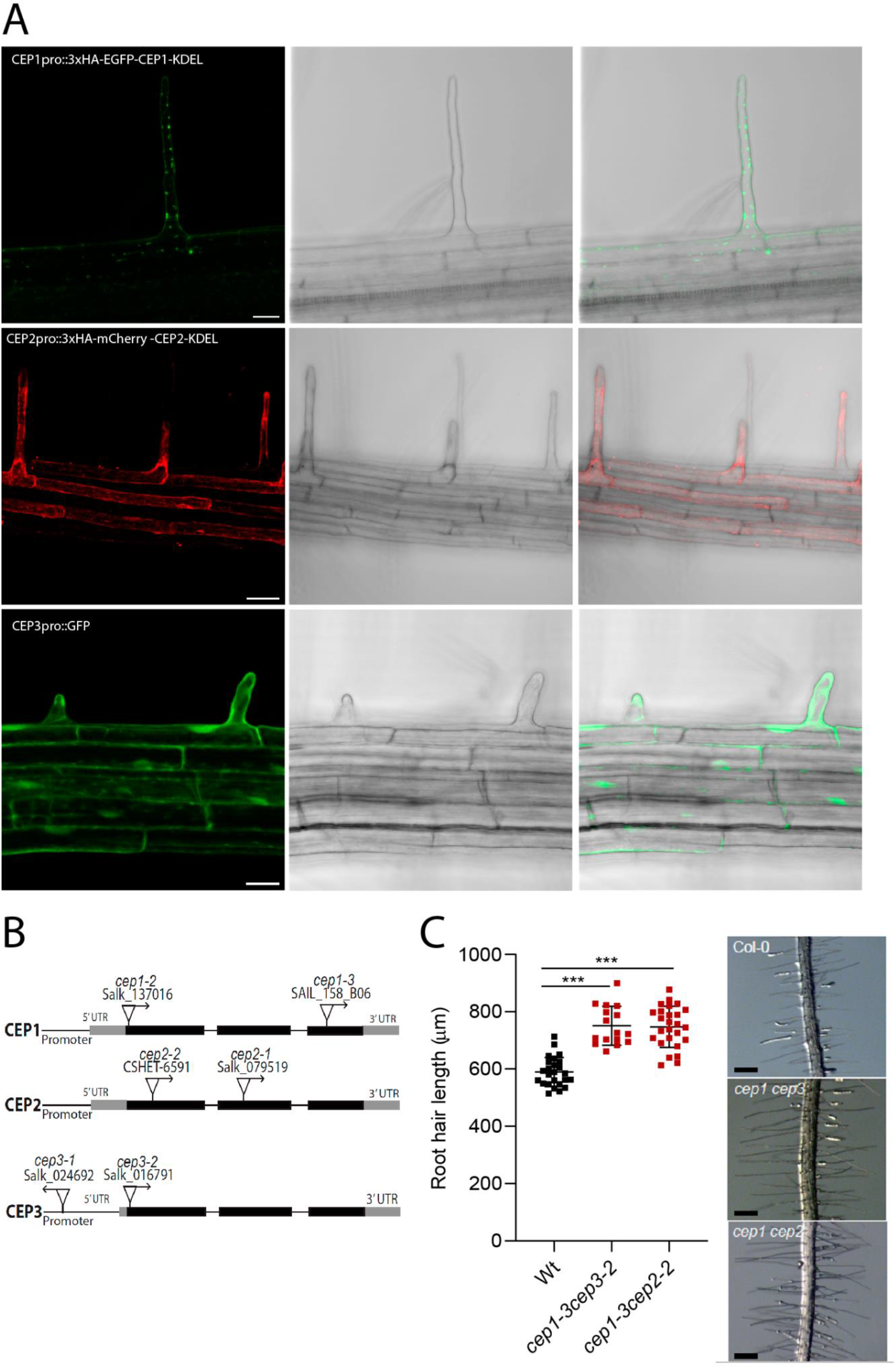
AtCEP1, AtCEP2 and AtCEP3 are expressed in root hairs and negatively regulate root hair growth. (**A**) AtCEP1 and AtCEP2 are expressed in root trichoblasts. Translational reporters for AtCEP1 (AtCEP1pro::PRE-PRO-GFP-KDEL) and AtCEP2 (AtCEP2pro::PRE-PRO-mCherry-AtCEP2-KDEL). AtCEP3 are expressed in both atricoblasts and tricoblasts. Transcriptional reporters of AtCEP3 (AtCEP3pro::GFP) in the root differentiation zone and specifically in RHs. Scale bar = 200 μm. (In the left) confocal microscopy image, (in the center) bright field microscopy image and (in the right) merge of two. (**B**) Scheme of AtCEP1, AtCEP2 and AtCEP3 genes showing introns (thin lines), exons (rectangles) and positions of T-DNA insertions. (**C**) Quantitative analysis of RH length (mean ± s.e.m., n= 200) in Wt Col-0 and *ceps* mutants. NS= not significant difference. Data are shown as the mean ± SEM, (n=20). Asterisks indicate significant differences from the Wt according to an ANOVA test with p<0.05. (On the right) Selected images of RHs in Wt and in single and double *cep* mutants. Scale bar = 200 μm.

To determine the nature of those organelles, we analysed for co-localization with established markers for different compartments (**Figure 2A**). The AtCEP2 labelled spotted and it did not co-localize or at very low levels with markers for the Golgi apparatus NAG1-GFP (0.10+-0.13) and N-ST-YFP (0.25+-0.16) (**Figure 2A**). However, we found higher co-localization levels in round or spindled shaped compartments with the ER membrane marker KKXX-GFP-KDEL (0.63+-0.16) and with HDEL-GFP lumenal ER marker (0.35+-0.15), indicating that AtCEP2 resides in an ER-derived compartment in root trichoblasts (**Figure 2B**). Indeed, AtCEP2 was observed in ER-derived compartments that resembled ricinosomes and ER-Bodies in leaf, hypocotyl, and root cap cells (Hierl et al. 2014; Howing et al. 2014). In addition, CEP2 mCherry was not detected in the apoplast space when plasmolysis was performed in root hairs (**Figure 2C**). On the other hand, apoplast targeting of AtCEPs has not been reported previously, possibly indicating that AtCEPs process their substrates within the secretory pathway. This confirms that AtCEP2 is primarily targeted to the ER compartment in root trichoblasts. We cannot rule out the possibility of a low level of an apoplast AtCEP2 expression. To analyse the expression of *AtCEP3*, we used a promoter fragment of 2-0Kb to drive GFP expression. *AtCEP3pro::GFP* fluorescence was also high in root trichoblasts, although not specific to this root epidermal cell-type (**Figure 1A-B**; **Figure S1**). However, the specific expression of AtCEP1 and AtCEP2 in root trichoblast cells and the expression of AtCEP3 in the epidermis indicates that these genes may play a role in growth of these specialized cells. To address if they are involved in RH polar growth we isolated T-DNA mutants for all three AtCEP genes (**Table S2** and **Figure 1C**; **Figure S2**). We characterized at least two T-DNA alleles for each *AtCEP* gene. Single mutants for *AtCEP1*, *AtCEP2* and *AtCEP3* showed similar phenotype to Wt Col-0 (**Figure S2**) while the double mutants *atcep1-3 atcep3-2* and *atcep1-3 atcep2-2* showed increased RH growth (up to 20% longer) when compared to Wt Col-0 (**Figure 1D**). These suggested that all three AtCEPs, AtCEP1-AtCEP3, redundantly restrict RH growth. Together, these results indicate that Arabidopsis AtCEP1-AtCEP3 proteins are expressed in RHs where they negatively regulate RH cell growth.

**Figure 2.**
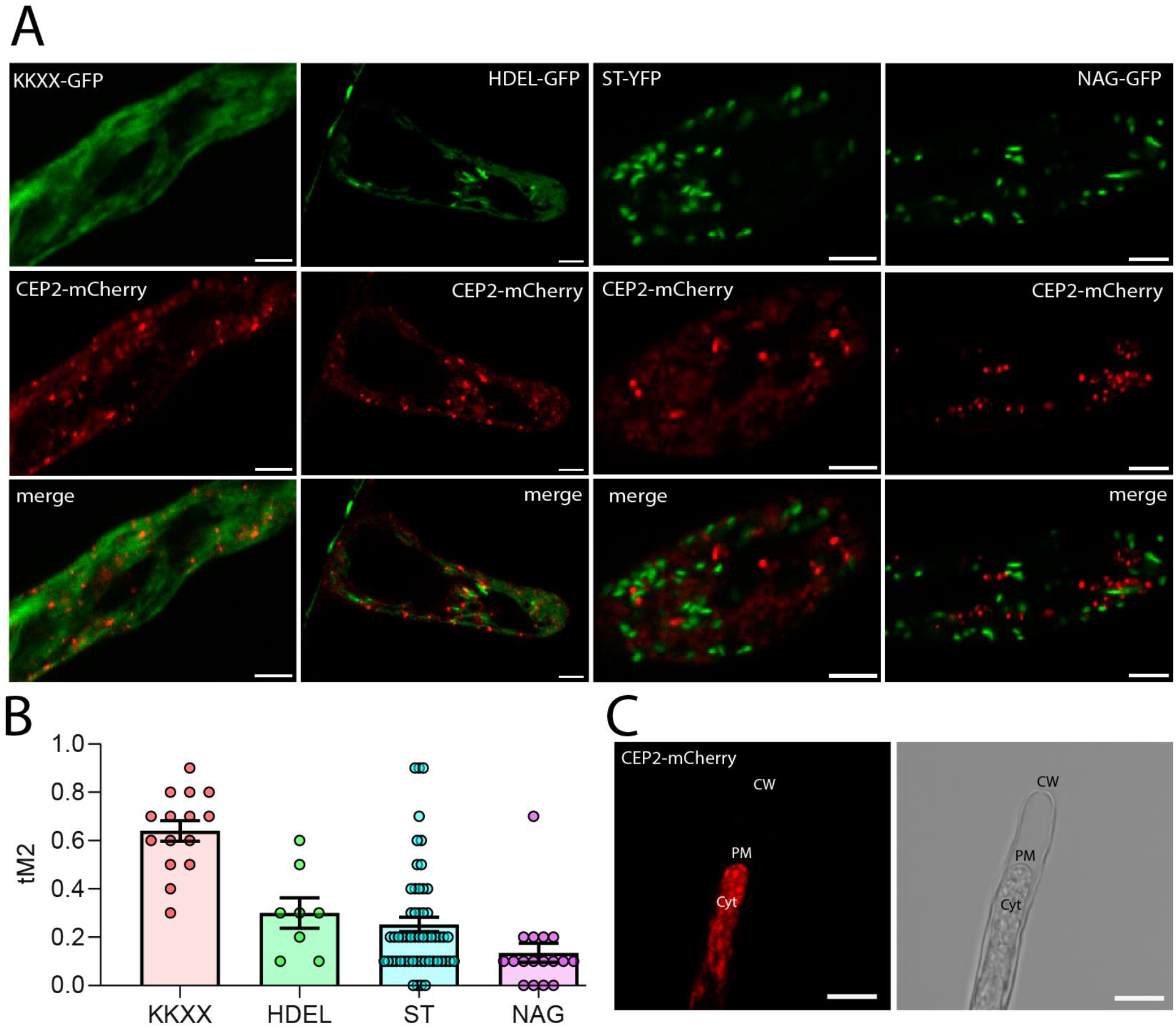
AtCEP2 colocalizes mostly with an ER-membrane marker and, to a lower extent, to Golgi markers in root hair cells. (**A**) Co-localization of AtCEP2-mCherry with markers for different subcellular compartments. Seven-day-old AtCEP2mCherry seedlings were grown (n= 10 roots with 1–5 root hair cells each). HDEL-GFP is a Lumen Endoplasmic reticulum (ER) marker and KKXX-GFP is an ER membrane marker. NAG-GFP is a cis-Golgi marker and ST-YFP is a trans-Golgi network marker. Scale bar 5 μm. (**B**) Quantification of colocalization using Manders correlation coefficient. (**C**) Plasmolyzed root hair to show the lack of signal in the apoplastic regions of AtCEP2-mCherry. Scale bar 20 μm. (CW) Cell Wall; (PM) Plasma membrane; (Cyt) Cytoplasm.

NAC1 was previously identified as a regulator of lateral root development and *de novo* root organogenesis (Xie et al., 2000; Chen et al., 2016). Since NAC1 was shown to control *AtCEP1* and *AtCEP2* expression during *de novo* root organogenesis (Chen et al. 2016), we tested if a similar regulation could be taking place in RHs. Indeed, *in silico* analysis (eFP browser and Root Cell Atlas) and characterization of a *NAC1pro::GFP* reporter line supports that *NAC1* is highly expressed in root trichoblasts, specifically in the later stages of their development (**Figure 3A-B**; **Figure S3**). Moreover, a *nac1-2* null mutant displays significantly longer RHs than Wt plants (**Figure 3B**). In addition, when we expressed *NAC1* fused to the repression domain SRDX (Hiratsu et al., 2003) to specifically suppress the expression of NAC1 target gene (35Spro::NAC1-SRDX-1 and 35Spro::NAC1-SRDX-2 lines) we observed an elongated RH phenotype (**Figure 3B**). Then, to analyse if *NAC1* overexpression would trigger a direct effects on RH growth possibly by upregulation of AtCEP1/AtCEP2, we used the pER8:3pro::×FLAG-NAC1 β-estradiol-inducible line (Zuo et al., 2000). In the presence of estradiol, an 75-fold induction of *NAC1* levels was observed (**Figure S4**). We then tested if the transcription factor NAC1 activates *AtCEP1-AtCEP3* expression by several folds in developing roots, as was shown for *de novo* adventitious root formation assay (Chen et al. 2016). The estradiol-inducible NAC1-FLAG line increased *CEP1* expression by 1.6 folds and *CEP2* by 4-folds while no changes were detected for *CEP3*. (**Figure S4**). On the contrary, the *nac1-2* mutant showed lower levels of transcripts of all three AtCEPs, AtCEP1-AtCEP3 (**Figure 3C**). Together, these results indicate that NAC1 activates the expression of *AtCEP1-AtCEP3*, and in consequence, controls RH growth, possibly through their EXT-processing activity. To determine if NAC1 controls the expression of *AtCEP* genes by direct binding to their promoter regions, we searched for putative NAC-binding sites in open chromatin regions, according to publicly available ATAC-Seq datasets (Maher et al., 2018). According to ChIP-qPCR using pER8:3pro::×FLAG-NAC1 plants (60x induction) and anti-FLAG antibodies, *AtCEP1*, *AtCEP2*, and lo a lower extent *AtCEP3*, all appeared as direct targets (**Figure 3D**), as revealed in comparison to the previously identified direct target *E2Fa* (Xie et al. 2023). All together, this confirms that NAC1 controls the expression of all three CEPs, *AtCEP1, AtCEP2,* and *AtCEP3* impacting on RH growth.

**Figure 3.**
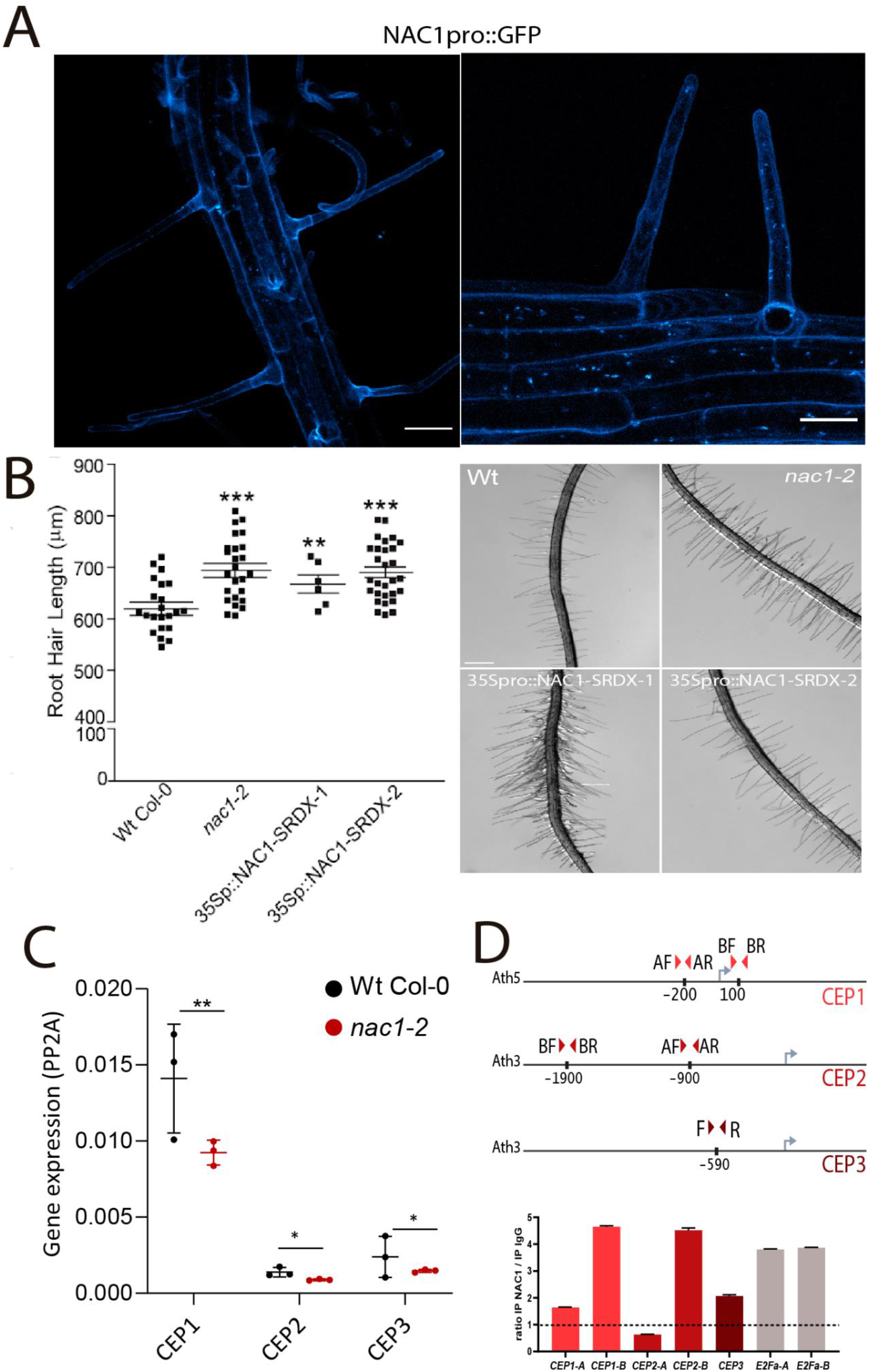
NAC1 is expressed and regulates root hair growth linked to AtCEP1-AtCEP3 expression. (**A**) NAC1pro:GFPexpression in roots and RHs. (**B**) RH phenotype in *nac1-2* mutant and two constitutive negative NAC1 lines. Quantitative analysis of RH length (mean ± s.e.m., n= 200). Data are shown as the mean ± SEM, (n=20). Asterisks indicate significant differences from the Wt Col-0 according to an ANOVA test with p<0.05. (**C**) qPCR analysis of AtCEP1, AtCEP2, and AtCEP3 in Wt Col-0 and *nac1-2* mutant. RNA was extracted from the roots of seedlings. *PP2A* was used as a control and amplification was performed for 30 cycles. Black arrowheads represent the region amplified by the primers used for the RT-PCR. Asterisks indicate significant differences from the Wt Col-0 according to an ANOVA test with (**) p<0.01 and (*) p<0.05. (**D**) ChIP-qPCR analysis of NAC1 binding to AtCEP1, AtCEP2 and AtCEP3 promoter regions. Schemes of the loci showing the location of the fragments analyzed by ChIP-qPCR are shown in the upper part. Primers were designed analyzing ATAC-seq experiments in regions where the chromatin is accessible (Maher et al., 2018). The enrichment was measured relative to the negative control ACTIN.

In previous studies, AtCEPs were shown to be involved in the processing of EXT proteins (Greenwood et al., 2005; Helm et al., 2008; Hierl et al., 2012). We thus tested if the reduction of AtCEP activity in the mutants had an effect on the EXTs secreted and insolubilized in the RH cell walls. To this end, we used an EXT-reporter carrying a tdTomato tag (SS-TOM-Long-EXT) that is resistant to the acidic pH, characteristic of cell wall apoplast (**Figure 4A**). Importantly, expression of the reporter did not affect the polar growth of RHs (Martinez Pacheco et al. 2022), making it an ideal probe for monitoring *in situ* alterations in the arrangement of cell wall EXTs. We measured the cell wall fluorescence signal from the SS-TOM-Long-EXT construct and its controls the SS-TOM construct in the apical zones of RHs under plasmolysis. Plasmolysis allowed us to retract the plasma membrane and detect the EXT-signal coming specifically from the cell walls. Interestingly, cell wall stabilization/insolubility of SS-TOM-Long-EXT in the RH tip was drastically reduced in *cep1 cep3* double mutant when compared to Wt Col-0 plants. To test if the total expression of the SS-TOM-Long-EXT construct was similar in both genetic backgrounds (Wt Col-0 vs *atcep1-3 atcep3-2* double mutant) without plasmolysis treatment, the overall signal was quantified (**Figure 4B**) and an overall higher signal was detected in *cep1 cep3* double mutant than in Wt Col-0 root hairs. These results suggest that the SS-TOM-Long-EXT reporter tested in the apical zone of the RHs is differentially modified by deficient AtCEPs activities possibly during the EXT processing in the secretory pathway.

**Figure 4.**
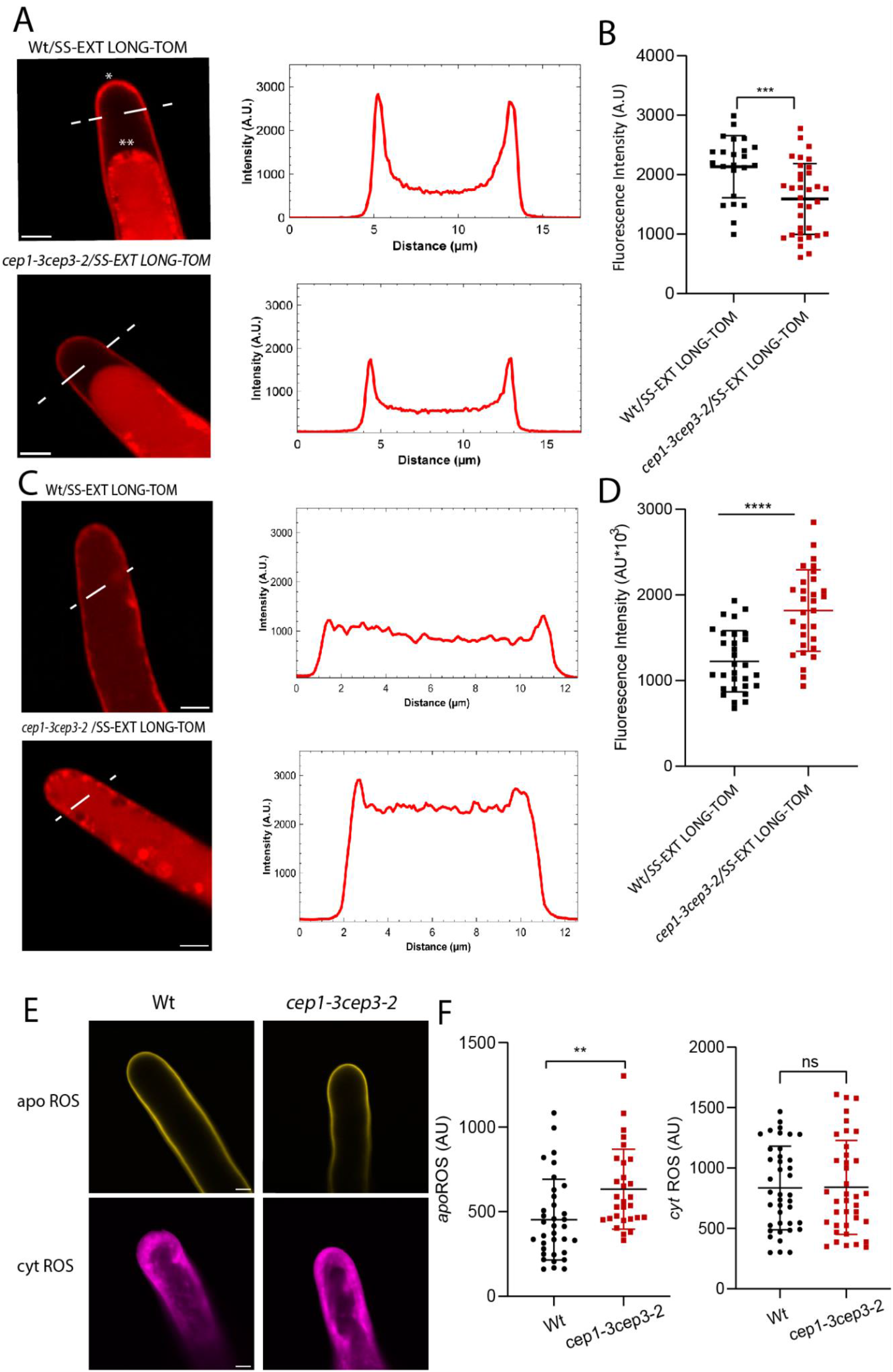
EXT secretion/insolubilization and apoplastic ROS are affected by AtCEP1-AtCEP3 in root hairs. (**A**) Signal of SS-EXT LONG-TOM in the apical zone of RHs in the *cep1-3 cep3-2* double mutant. Cells were plasmolyzed with a mannitol 8% solution. (On the left) Intensity profiles across a dotted line in the RH tips. (**B**) Each point in the graph is the signal derived from a single RH tip. Fluorescence AU data are the mean ± SD SD (N = 3), two-way ANOVA followed by a Tukey–Kramer test; (***) p<0.01. Results are representative of two independent experiments. NS = non-significant differences. Scale bars = 5 μm. (**C**) Signal of SS-EXT LONG-TOM in the apical zone of RHs in the *cep1-3 cep3-2* double mutant. (On the left) Intensity profiles across a dotted line in the RH tips. (**D**) Each point in the graph is the signal derived from a single RH tip. Fluorescence AU data are the mean ± SD SD (N = 3), two-way ANOVA followed by a Tukey– Kramer test; (****) p<0.001. Results are representative of two independent experiments. NS = non-significant differences. Scale bars = 5 μm. (**E**) Cytoplasmic ROS (_cyt_ROS) levels were measured using the hydrogen peroxide-selective dye Peroxy-Orange 1 (PO1) in apical areas of RHs in wild-type (Columbia Col-0) and in the double mutant *cep1-3 cep3-2*. Each point is the signal derived from a single RH tip. Fluorescence AU data are the mean ± SD (N= 20 RHs), two-way ANOVA followed by a Tukey–Kramer test; (**) p < 0.01. Results are representative of two independent experiments. Asterisks indicate significant differences. NS = non-significant differences. Scale bars = 2 μm. (**F**) Apoplastic ROS (_apo_ROS) levels were measured with Amplex™ UltraRed in apical areas of RHs in wild-type (Columbia Col-0) and in the double mutant *cep1-3 cep3-2*. Each point in (**F**) is the signal derived from a single RH tip. Fluorescence AU data are the mean ± SD (N =3), two-way ANOVA followed by a Tukey–Kramer test. Results are representative of two independent experiments. Scale bars = 2 μm.

Since EXTs insolubilization in the cell walls of growing RHs is regulated by several factors including the Reactive Oxygen Species (ROS) homeostasis (Martinez Pacheco et al. 2022; Marzol et al. 2022), we tested if the effect of these AtCEPs might affect global ROS levels. We measured total cytoplasmic ROS (_cyt_H_2_O_2_) with the hydrogen peroxide-selective dye Peroxy-Orange 1 (PO1), as a permeable boronate-based dye that is non-fluorescent in its reduced form, but becomes fluorescent when irreversibly oxidized by H_2_O_2_ (Dickinson et al., 2010). Apoplastic ROS (_apo_H_2_O_2_) levels were determined in the RH tips with the cell-impermeable Amplex™ UltraRed Reagent (Mangano et al. 2016; Matinez-Pacheco et al. 2022; Marzol et al. 2022) (**Figure 4B**). The *atcep1-3 atcep3-2* showed higher levels of _apo_H_2_O_2_ in RH tips compared to Col-0, whereas the _cyt_H_2_O_2_ were similar in both genotypes. This indicates that ROS homeostasis is also changed in the *ceps* mutant background, which can also affect the EXTs insolubilization in the cell walls. Changes in ROS homeostasis in apoplastic and cytoplasmic apical areas were also detected in overgrowing root hairs when apoplastic type-III peroxidases (PRXs) related to EXTs were overexpressed (Marzol et al. 2022; Pacheco et al. 2022). This may imply a close relationship between the proper status of processed EXTs in the cell walls, ROS homeostasis in the apical zone and balanced root hair growth as previously hypothesised (Mangano et al. 2018).

Finally, to test if AtCEP1-AtCEP3 might be able to interact with single-chain EXTs, we performed homology modelling with one cysteine endopeptidase of RcCysEP and a cysteine protease from *Ambrosia artemisiifolia* (pdbs 1S4V and 5EF4 respectively) known to be able to cleave and process EXT-like substrates under *in vitro* conditions. AtCEP1-AtCEP3 proteins share a sequence identity of 66-74% with the most well-characterized *Ricinus communis* RcCysEP (**Figure S5**). By *docking* analysis, we obtained interaction energies (Kcal/mol) for all three AtCEPs proteins and they were compared to RcCysEP 1SV4. We analyzed docking with four different short EXT peptides from non-hydroxylated to fully *O*-glycosylated peptide (Velasquez et al. 2015; Marzol et al. 2022) (**Figure S6**). It was previously shown that mutants carrying under-*O*-glycosylated EXTs and related EXT-proteins have severe defects in RH growth (Velasquez et al. 2011; Velasquez et al. 2015). Our docking results for these three AtCEPs showed consistent interaction energy differences that depend on the EXT glycosylation state, being higher for non-*O*-glycosylated species. In general, we observed higher interaction energies (higher negative values) for (non)-hydroxylated EXT species than for *O*-glycosylated EXT variants. When we compared interaction energies among different AtCEPs interacting with EXT substrates, we observed that AtCEP1 and AtCEP3 displayed the highest interaction activity with the (non)-hydroxylated EXT species (**Figure S6**). Overall, this likely indicates that AtCEP1 and AtCEP3 might interact with EXT substrates and possibly catalyse proteolysis in open regions of the EXT backbones with little or no *O*-glycosylation while CEP2 might prefer *O*-Glycosylated regions. This is in agreement with previous studies suggesting that high levels of *O*-glycosylation in certain proteins physically restrict its degradation or processing.

## Conclusions

It was previously considered that up-regulation of AtCEPs was related to the processing of EXT since EXT promotes wound healing, and this might be a barrier for the emergence of regenerated root tips. Therefore, it was hypothesized that the NAC1-AtCEPs antagonises EXT-mediated wound healing, and this allows the emergence of regenerated root tips (Chen et al. 2016). However the overall results in our study suggest that NAC1 directly activates *AtCEP1*, *AtCEP2* and *AtCEP3* expression, which in turn represses polar-cell expansion in growing RHs. By a colocalization analysis with several subcellular markers, we detected AtCEP2 in ER-derived compartments but not in the apoplast. Based on our results, it is plausible that AtCEP1-AtCEP3 could act together as components on the EXT and EXT-related protein quality control program and proper EXTs protein processing. So far there is no information regarding the existence of a quality control pathway for *O*-glycosylated proteins in the same manner to the well-known and conserved CNX (calnexin)/calreticulin lectins linked to ERAD (ER-associated degradation)-program (e.g. with OS9, HRD3/SEL1L components) as it happens for misfolded *N*-glycosylated proteins (Su et al. 2011; Su et al. 2012; Strasser 2016). It is also unknown how the *O*-glycosylation machinery required for a proper processing of EXTs and EXT-related proteins in the secretory pathway (Velásquez et al. 2012; Marzol et al. 2018) are coordinated and controlled. Our findings pave the way for the discovery of novel functions for AtCEPs in cells that are not undergoing PCD but are characterized by extensive cell wall expansion or remodelling, possibly as a result of EXTs cleavage. The involvement of a protease in cell wall remodelling indicates the significance of the protein component of the cell wall in this process. Insights into the underlying mechanisms, such as the membrane fusion of ricinosomes with the plasma membrane during root development, similar to other ER-derived vesicles, will be intriguing hypotheses to investigate. Recently, a vacuolar processing enzyme (βVPE) was found to control the maturation of AtCEP1 transforming this pro-protease into mature protease while in *vpe* mutants, the maturation of AtCEP1 and other proteases was severely inhibited (Cheng et al. 2020). It is still unclear if VPE may be also acting in root trichoblasts to control AtCEPs processing. Root trichoblasts provide an excellent model system to dissect the molecular components of the AtCEPs-EXT *O*-glycosylation pathway required for polar-growth. Finally, in our work we found that misregulation of EXT processing by CEPs enhances root hair growth while previously we found that enhancing the expression of type-III apoplastic Peroxidases PRX44/PRX73 (Marzol 2023) and PRX62 (Pacheco et al. 2022) also ends up in a similar root hair phenotype. Overall, it seems that subtle changes in EXTs processing and crosslinking has a direct effect on root hair growth as previously demonstrated for the impact of proline-hydroxylation and *O*-Glycosylation status of EXTs in plant cell growth (Velasquez et al. 2011, 2015). Currently, one of the main obstacles in the study of these complex glycoproteins is to visualise the chemical changes occurring in EXTs (proline hydroxylation, *O*-glycosylation, oligomerization and tyr-crosslinking) while they are being synthetized and then secreted *in situ* in the cell walls in an *in vivo* conditions.

## Experimental Procedures

### Plant and Growth Conditions

*Arabidopsis thaliana* Columbia-0 (Col-0) was used as the Wt genotype in all experiments. All mutants and transgenic lines tested are in this ecotype. Seedlings were germinated on agar plates in a Percival incubator at 22°C in a growth room with 16h light/8h dark cycles for 7-10 days at 140 μmol m^−2^s^−1^ light intensity. Plants were transferred to soil for growth under the same conditions as previously described at 22°C. For identification of T-DNA knockout lines, genomic DNA was extracted from rosette leaves. Confirmation by PCR of a single and multiple T-DNA insertions in the target AtCEP and NAC genes were performed using an insertion-specific LBb1 or LBb1.3 (for SALK lines) or Lb3 (for SAIL lines) primer in addition to one gene-specific primer. To ensure gene disruptions, PCR was also run using two gene-specific primers, expecting bands corresponding to fragments larger than in Wt. In this way, we isolated homozygous lines (for all the genes mentioned above). Mutant list is detailed in **Supplementary Table S1**. Lines expressing AtCEP2pro::pre-pro-3xHA-mCherry-AtCEP2 were crossed with ER and Golgi marker lines to obtain the double transgenic AtCEP2 + organelle marker.

### Root Hair Phenotype

Seeds were surface sterilised and stratified in darkness for 3 d at 4°C and they were grown *in vitro* on a specific condition and medium in a plant growth chamber in continuous light (120 mol s-1 m-2) at 22°C. The quantitative analyses of RH phenotypes of Col-0 and transgenic lines were made the last day of the growth conditions described in the two previous sections. Images were captured using an Olympus SZX7 Zoom Stereo Microscope (Olympus, Tokyo, Japan) equipped with a Q-Colors digital camera and Q CAPTURE PRO 7 software (Olympus). Results were expressed as the mean SD using the GRAPHPAD PRISM 8.0.1 (GraphPad Software, Boston, MA, USA) statistical analysis software. Results are representative of three independent experiments, each involving 15–20 roots.

### AtCEP imaging

Root hairs were ratio imaged with the Zeiss LSM 510 laser scanning confocal microscope (Carl Zeiss) using a 40X oil-immersion, 1.2 numerical aperture objective. EGFP (473– 505nm) and mCherry (526–536 nm) emissions were collected using a 458-nm primary dichroic mirror and the Meta-detector of the microscope. Bright-field images were acquired simultaneously using the transmission detector of the microscope. For time-lapse analysis, images were collected every 3 or 5 s. Image sequences were analyzed using the Template Matching and Slice Alignment plug-in for ImageJ. Fluorescence intensity was measured in 7 µm ROI at the RH apex.

### Transcriptional reporter generation

Vectors based on the Gateway cloning technology (Invitrogen) were used for all manipulations. Promoter regions (2Kb) were PCR-amplified with AttB recombination sites. PCR products were first recombined in pDONOR207 and transferred into pMDC111. Transgenic lines used in this study are described in **Table S2**.

### AtCEPs expression

To quantify AtCEP1 and AtCEP2 expression levels, lines AtCEP1pro::3xHA-EGFP AtCEP2pro::3xHA-mCherry were used. The fluorescence intensity of GFP and mCherry was measured respectively. All RHs growth stages were measured. Ten days old seedlings were removed from the medium and placed in a slide containing a drop of liquid MS 0.5X in the dark, and images were obtained using a Zeiss Imager A2 epifluorescence microscope. The objective used was 10X, 0.3 numerical aperture, exposure time 2 seconds. The lasers used were suitable for each fluorophore: GFP (GFP λ max ex. = 488 nm, λ max em. = 507 nm) and mCherry (λ max ex. = 536 nm, λ max em. = 632 nm) The Wt Col-0 line was used to rule out autofluorescence noise. The images were analyzed using ImageJ 1.50b software. To measure fluorescence intensity levels (represented in arbitrary units: UA), both a circular region of interest within the RH apex (ROI) and the total area of RH were selected. All RHs from 6 seedlings were analyzed. The reported values are the mean ± standard error (mean ± SEM).

### Subcellular localization of AtCEP2

Lines expressing CEP2pro:pre-pro-3xHA-mCherry-CEP2 were crossed with ER and Golgi marker lines to obtain the double transgenic CEP2 + organelle marker. Co-localization analysis was performed using the BIOP implementation (Battistella et al. 2019) of the JACoP plugin (Bolte and Cordelières 2006) in ImageJ (1.53t) on individual root hairs for which ROIs (Region Of Interest) were drawn manually. Background intensity was subtracted from each channel as the mean intensity of the autofluorescence control + 2SD. Different marker expression levels excluded the usage of fixed cutoff values for all images, so the histogram-derived Otsu method was chosen based on visual inspection of the thresholded images for both channels. To compare the relative localization of CEP2pro:pre-pro-3xHA-mCherry-CEP2 (Ch2) in each of the different subcellular compartments (Ch1), we used Mander’s M2 overlap coefficient (Manders et al. 1993) to measure the proportion of positive pixels in Ch2 that co-occur with positive pixels in Ch1. Confocal microscopy images were acquired on a Zeiss LSM710 microscope with an EC Plan-Neofluar 40x/1.30 oil objective. Images were acquired sequentially on different tracks in order to avoid excitation and emission bleed-through using the following emission ranges for the individual channels (8-bit pixel depth): Ch1 519−589 nm, Ch2 594-690 nm. Pixel size was set at 100 nm following Nyquist sampling criterion and pinhole was adjusted to obtain a 3.6 µm thick optical slice. For each experimental replicate, 15-25 total individual root hairs from five different plants per subcellular compartment marker were imaged.

### Chromatin immunoprecipitation Assay

Chromatin immunoprecipitation (ChIP) assays were performed on pER8:3xFLAG-NAC1 plants (Chen et.al 2016, DOI: 10.1104/pp.15.01733) mainly as described in Ariel et al. (2020). Plants were grown for 10 days in plates containing MS 0,5X medium (pH 5,7; 0.8% agar) placed vertically in a culture chamber at 22°C and continuous light (140μmol/m2.sec). After 10 days, the plates were placed horizontally and treated with β-estradiol 10 µM solution for 3h. The expression of NAC1 was checked by qPCR (supplementary table Sx). Chromatin was cross-linked with formaldehyde 1% for 10 min at room temperature. Cross-linking was stopped by adding glycine (125 mM final concentration) and incubating for 10 min at room temperature. Crosslinked chromatin was extracted by cell resuspension, centrifugation, cell membrane lysis, and sucrose gradient as previously described (Ariel et al., 2020). Nuclei were resuspended in Nuclei Lysis Buffer and chromatin was sonicated using a water bath Bioruptor Pico (Diagenode; 30 s on / 30 s off pulses, at high intensity for 10 cycles). Chromatin samples were incubated for 12 h at 4 °C with Protein G Dynabeads (Invitrogen) precoated with the antibodies anti-FLAG (Abcam ab18230) or anti-IgG (Abcam ab6702) as a negative control. Immunoprecipitated DNA was recovered using Phenol:Chloroform:Isoamilic Acid (25:24:1; Sigma) and analyzed by qPCR using the primers listed in **Supplementary Table S3**. Two regions upstream of the E2Fa gene were used as a positive control (Xie and Ding, 2022). Untreated sonicated chromatin was processed in parallel and considered the input sample. The GraphPad Prism 6 software was used to analyze the data and produce the graphs.

### Apoplastic and cytoplasmic ROS measurements

To measure ROS levels in root trichoblasts, 8 days-old Arabidopsis seedlings grown in continuous light were used. For cytoplasmic ROS, the seedlings were incubated in darkness for 10 min with H2O2 and were visualized with Peroxy-Orange 1 (PO1). PO1 was dissolved in DMSO to make a 500 μM stock and was further diluted in water to make a 50 μM working solution. Seedlings were incubated in PO1 for 15 min in the dark and were then rinsed with water and mounted in water for imaging and observed with Zeiss Imager A2 Epifluorescence Microscope (Zeiss, Germany) (Plain Apochromat 40X/1.2 WI objective, exposure time 25 ms). Images were analyzed using ImageJ software. To measure ROS levels, a circular region of interest was chosen in the zone of the RH tip cytoplasm. To measure apoplastic ROS, the seedlings were incubated with 50 μM Amplex™ UltraRed Reagent (AUR) (Molecular Probes, Invitrogen) for 15 min in darkness and rinsed with liquid 0.5X MS media (Duchefa, Netherlands). Root hairs were imaged with a Zeiss LSM5 Pascal (Zeiss, Germany)) laser scanning confocal microscope (Excitation 543 nm argon laser; Emission: 560–610 nm, Plain Apochromat 40X/1.2 WI objective). Quantification of the AUR probing fluorescence signal was restricted to apoplastic spaces at the RH tip and quantified using the ImageJ software. Fluorescence AU were expressed as the mean ± SD using the GraphPad Prism 8.0.1 (USA) statistical analysis software. Results of both ROS measurements are representative of two independent experiments, each involving 10–15 roots and approximately, between 10-20 RHs per root were observed.

### Treatments with *β*-Estradiol

*β-*Estradiol (Sigma-Aldrich,) was added to 0.5X MS media at a final concentration of 10 μM. The line ER8pro::3×FLAG-NAC1 was grown at 22°C for 4 days + 3 days of *β-* Estradiol induction. Root hairs phenotype was then quantified as indicated before.

### Quantitative reverse transcriptase PCR (qRT-PCR)

Total RNA was isolated from 10-d-old seedling roots (40 for each line) using the RNeasy Plant Mini Kit (Qiagen, Germany). cDNA was synthesized using TOPscript^TM^ RT DryMIX (dT18, Enzynomics, Korea). qRT-PCR analyses were performed using TOPrealTM qPCR 2x PreMIX (SYBR Green, Enzynomics, Korea) and Chromo4™ Four-Color Real-Time Detector (Bio-Rad, USA). Gene-specific signals were normalized relatively to *PP2A* (At1G69960) signals. Each qRT-PCR reaction was performed in triplicate, and each experiment was repeated three times using independent preparations of RNA. Primers used are as listed in **Table S3**.

### SS-TOM and SS-TOM-Long-EXT constructs

These construct were described in detil in Martinez Pacheco et al. 2022. Briefly, The secretory sequence (SS) from tomato polygalacturonase is MVIQRNSILLLIIIFASSISTCRSGT (2.8 kDa) and the EXT-Long sequence is BAAAAAAACTLPSLKNFTFSKNIFESMDETCRPSESKQVKIDGNENCLGGRSEQRTEKECFPVVSKPVDCSK GHCGVSREGQSPKDPPKTVTPPKPSTPTTPKPNPSPPPPKTLPPPPKTSPPPPVHSPPPPPVASPPPPVHSPPPPVASPPPPVHSPPPPPVASPPPPVHSPPPPVASPPPPVHSPPPPVHSPPPPVASPPPPVHSPPPPVHSPPPPVHSPPPPVHSPPPPVHSPPPPVASPPPPVHSPPPPVHSPPPPVHSPPPPVASPPPPVHSPPPPPPVASPPPPVHSPPPPVASPPPPVHSPPPPVASPPPPVHSPPPPVHSPPPPVHSPPPPVASPPPALVFSPPPPVHSPPPPAPVMSPPPPTFEDALPPTLGSLYASPPPPIFQGY* 395–(39.9 kDa). The predicted molecular size for SS-TOM protein is 54.2 kDa and for SS-TOM-EXT-Long Mw is 97.4 kDa.

### Modelling and molecular docking between AtCEP1-AtCEP3 and EXTs

Modelling and molecular docking: cDNA sequences of AtCEPs were retrieved from TAIR (AtCEP1: AT1G05240, AtCEP2: AT3G50990, AtCEP3: AT4G26010) and NCBI Nucleotide DB. Homology modelling was performed using an AtCEP from *Ricinus communis* and from *Ambrosia artemisiifolia* using modeller 9.14 (Sali et al. 1993), using the crystal structures 1S4V and 5EF4 as templates, available at the protein data bank. 100 structures were generated for each protein and the best scoring one (according to DOPE score) was picked. The *receptor* for the docking runs was generated by the prepare_receptor4 script from the autodock suite, adding hydrogens and constructing bonds. Peptides based on the sequence PYYSPSPKVYYPPPSSYVYPPPPS were used, replacing proline by hydroxyproline, and/or adding *O*-Hyp glycosylation with up to four arabinoses per hydroxyproline in the fully glycosylated peptide and a galactose on the serine, as it is usual in plant *O*-Hyp (Strasser 2016). Ligand starting structure was generated as the most stable structure by molecular dynamics (Velasquez et al. 2015a). All ligand bonds were set to be able to rotate. Docking was performed in two steps, using Autodock vina (Trott et al. 2010). First, an exploratory search over the whole protein surface (exhaustiveness 4) was done, followed by a more exhaustive one (exhaustiveness 8), reducing the search space to a 75×75×75 box centered over the most frequent binding site found in the former run.

## Acknowledgements

We thank ABRC (Ohio State University) for providing T-DNA lines seed lines and Gietl for providing some of the materials used in this study. J.M.E. is an investigator of the National Research Council (CONICET) from Argentina. This work was supported by grants from ANPCyT (PICT2019-00015 and PICT2021-0514 to J.M.E and PICT2018-00577 to E.M.) and from the Spanish Ministry of Science and Innovation MCIN/AEI/10.13039/501100011033 and FEDER “una manera de hacer Europa” (PID2021-128078NB-I00 to E.R.). In addition, this research is also funded by ANID – Programa Iniciativa Científica Milenio ICN17_022, NCN2021_010 and Fondo Nacional de Desarrollo Científico y Tecnológico [1200010] to J.M.E. The authors gratefully acknowledge the Microscopy and Imaging Facility at the Leloir Institute Foundation (FIL) for their support and assistance in the present work.

## Author Contribution

D.R.G performed all the experiments and analyzed the data. E.M isolated *cep* mutants and analyzed the data. Y.d.C.R. performed ROS experiments. A.H.R and E.A.M. carried out co-localization studies. A.A.A. and A.D.N. carried out the molecular modeling of AtCEPs-EXTs. L.F and F.A. carried out the ChIP experiment and analyzed the data. J.M. P., M.C., V.B.G., L.E.L, G.D.D. and C.B. analyzed the data. L.X., E.R and J.J.S.S. provided materials, analyzed the co-localization data and wrote the paper. E.R. wrote the paper. J.M.E. designed research, analyzed the data, supervised the project, and wrote the paper. All authors commented on the results and the manuscript. This manuscript has not been published and is not under consideration for publication elsewhere. All the authors have read the manuscript and have approved this submission.

## Supplementary Materials

Figure S1-S6

Tables S1-S3

**Figure S1.**
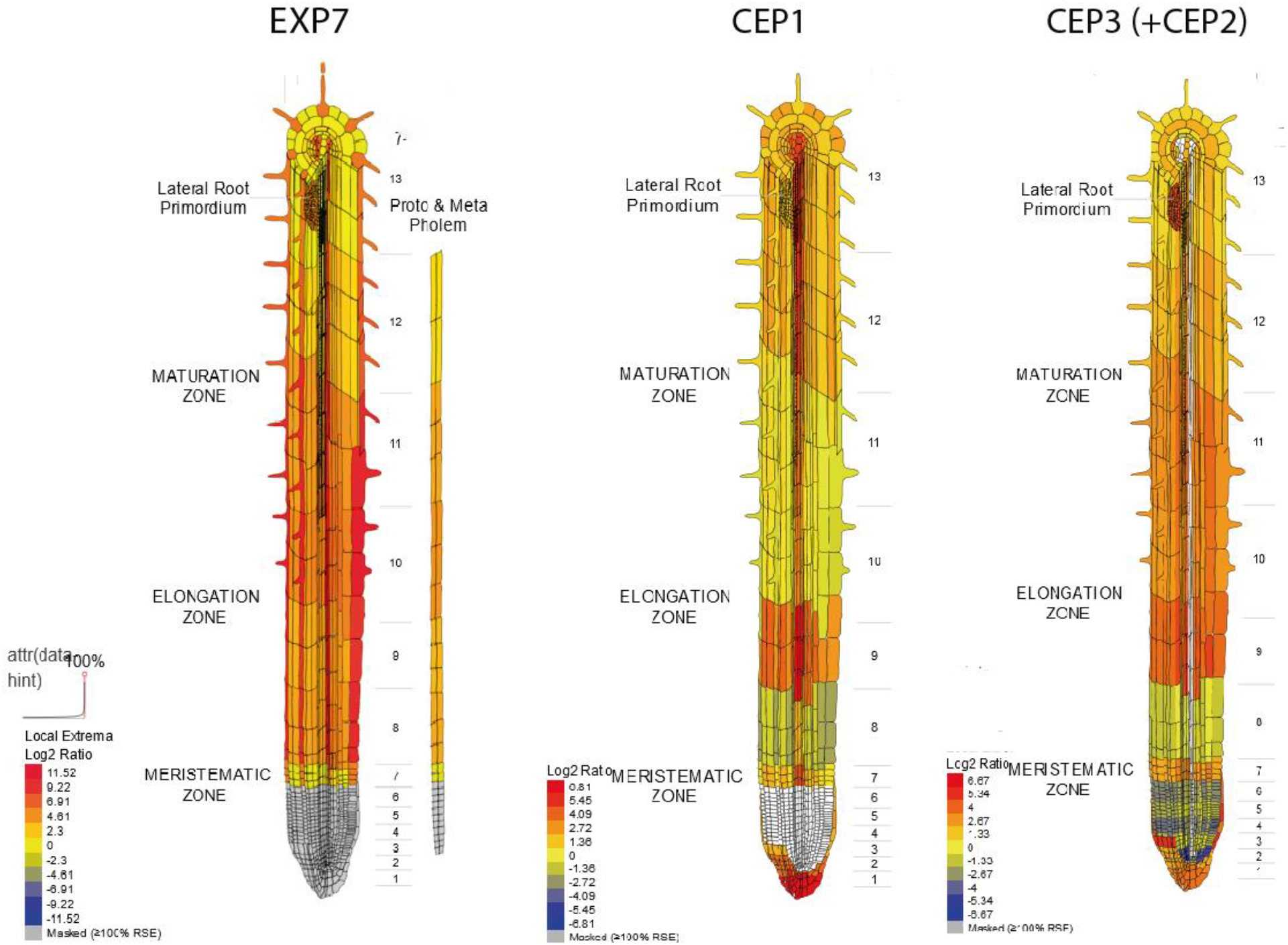
The *in silico* analysis of AtCEPs gene expression using Tissue Specific Root eFP (http://bar.utoronto.ca/eplant/). EXP7 was included as a root hair cell marker.

**Figure S2.**
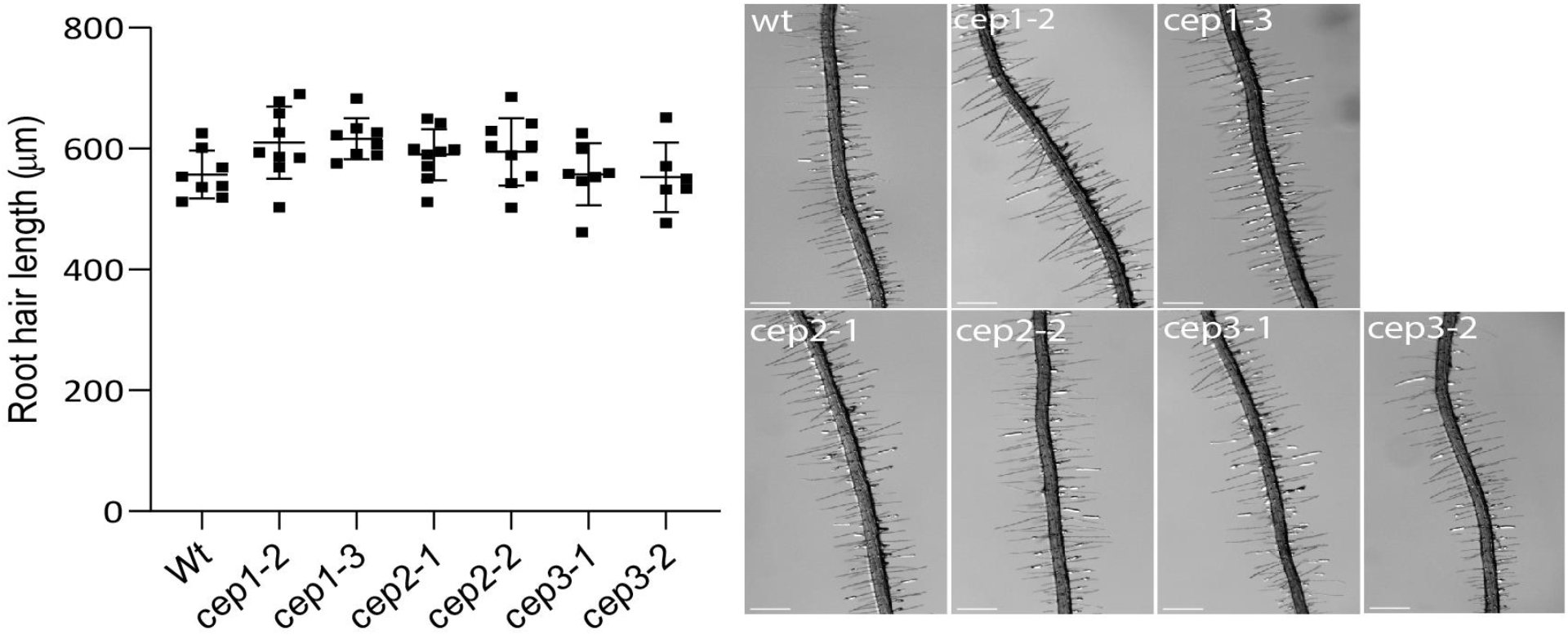
Single *cep* mutants do not perturb root hair growth. Quantitative analysis of RH length (mean ± s.e.m., n= 200) in Wt Col-0 and in single *cep* mutants. Data are shown as the mean ± SEM, (n=20). (On the right) Selected images of RHs in Wt and in single and double *cep* mutants. Scale bar = 200 μm.

**Figure S3.**
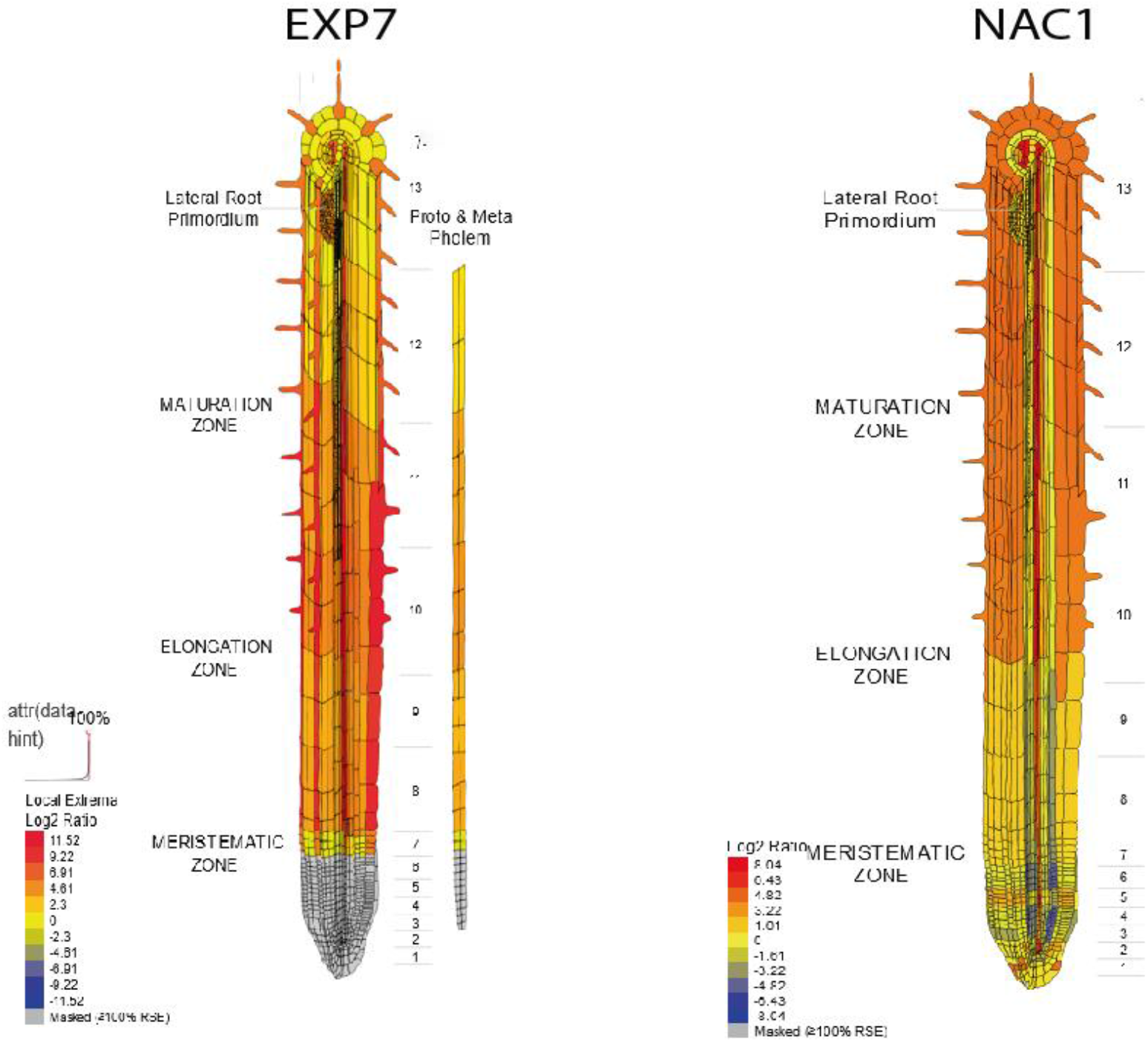
The *in silico* analysis of NAC1 expression using Tissue Specific Root eFP (http://bar.utoronto.ca/eplant/). EXP7 was included as a root hair cell marker.

**Figure S4.**
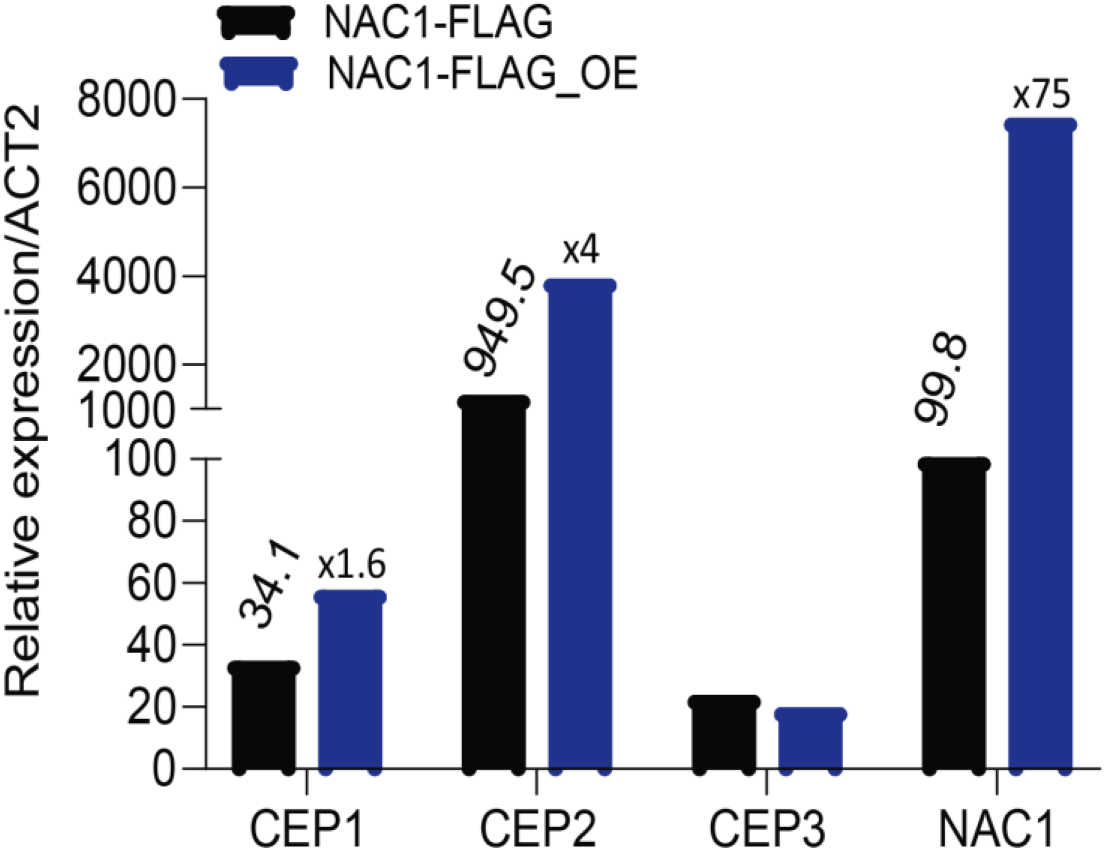
Induction of CEP1 and CEP2 by NAC1-FLAG inducible line (PER8pro:3×FLAG-NAC1). Gene-specific signals were normalized relatively to ACT2.

**Figure S5.**
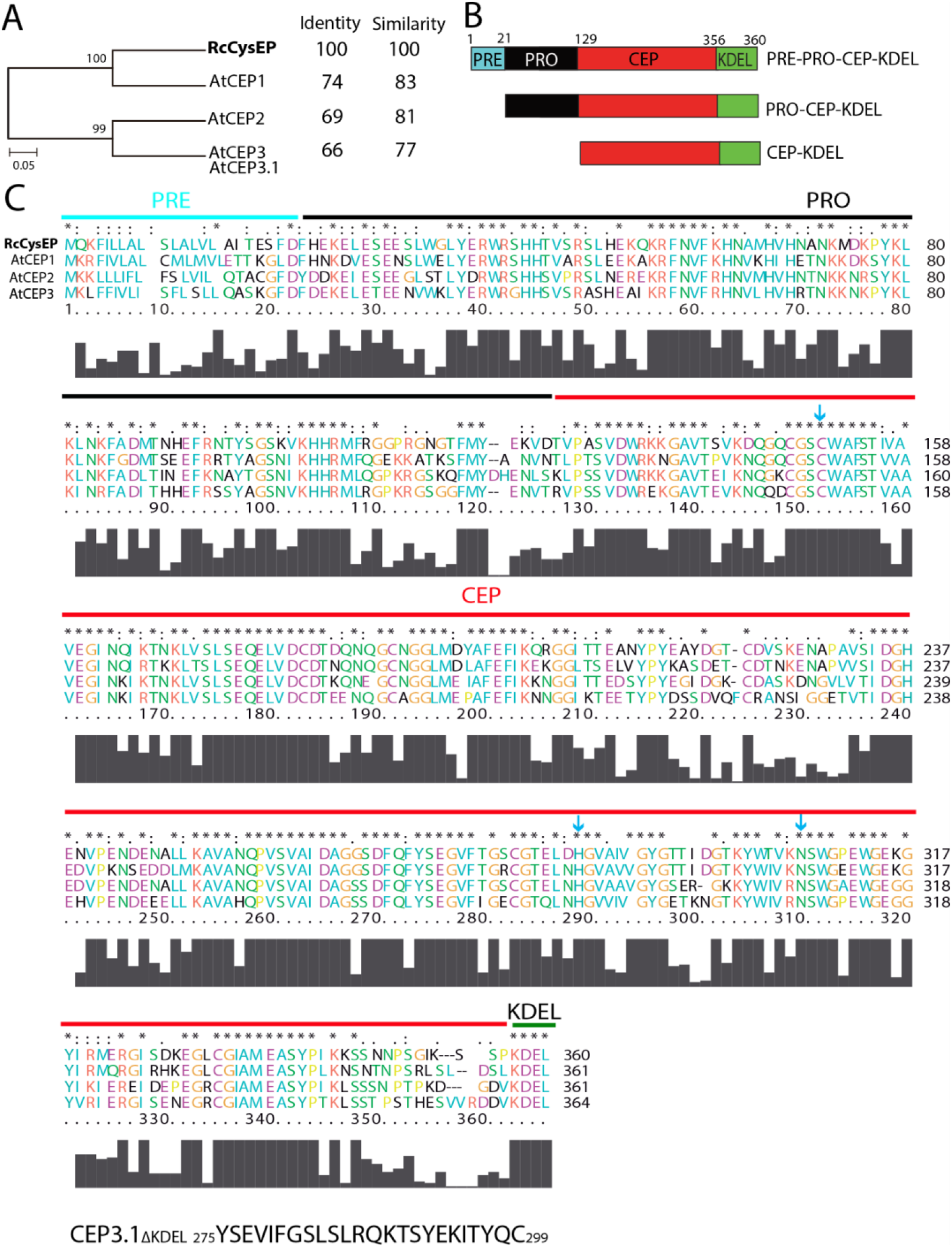
Protein alignment and domains of AtCEP1-AtCEP3 sequences from *Arabidopsis thaliana* with Ricinus CEP 1SV4 (RcCysEP). (A) Phylogenetic tree of Arabidopsis AtCEPs and Ricinus AtCEP. The phylogenetic analysis was carried out with MEGA6 (Tamura et al., 2007) using the Maximum Similarity method (Maximum Likelihood) (Saitou and Nei, 1987). The numbers in the nodes indicate the bootstrap values obtained for 1000 iterations. Scale represents the evolutionary distance, expressed as the number of substitutions per amino acid. (B) AtCEP protein domains. AtCEPs are synthesized as pre-pro-enzyme, which is then co-translationally synthesized into the ER, where the pre-sequence is removed, and pro-enzyme is finally released from the ricinosomes. KDEL is an ER retention signal peptide. (C) AtCEP1-AtCEP3 Protein alignment. PRE, PRO and KDEL domains are indicated.

**Figure S6.**
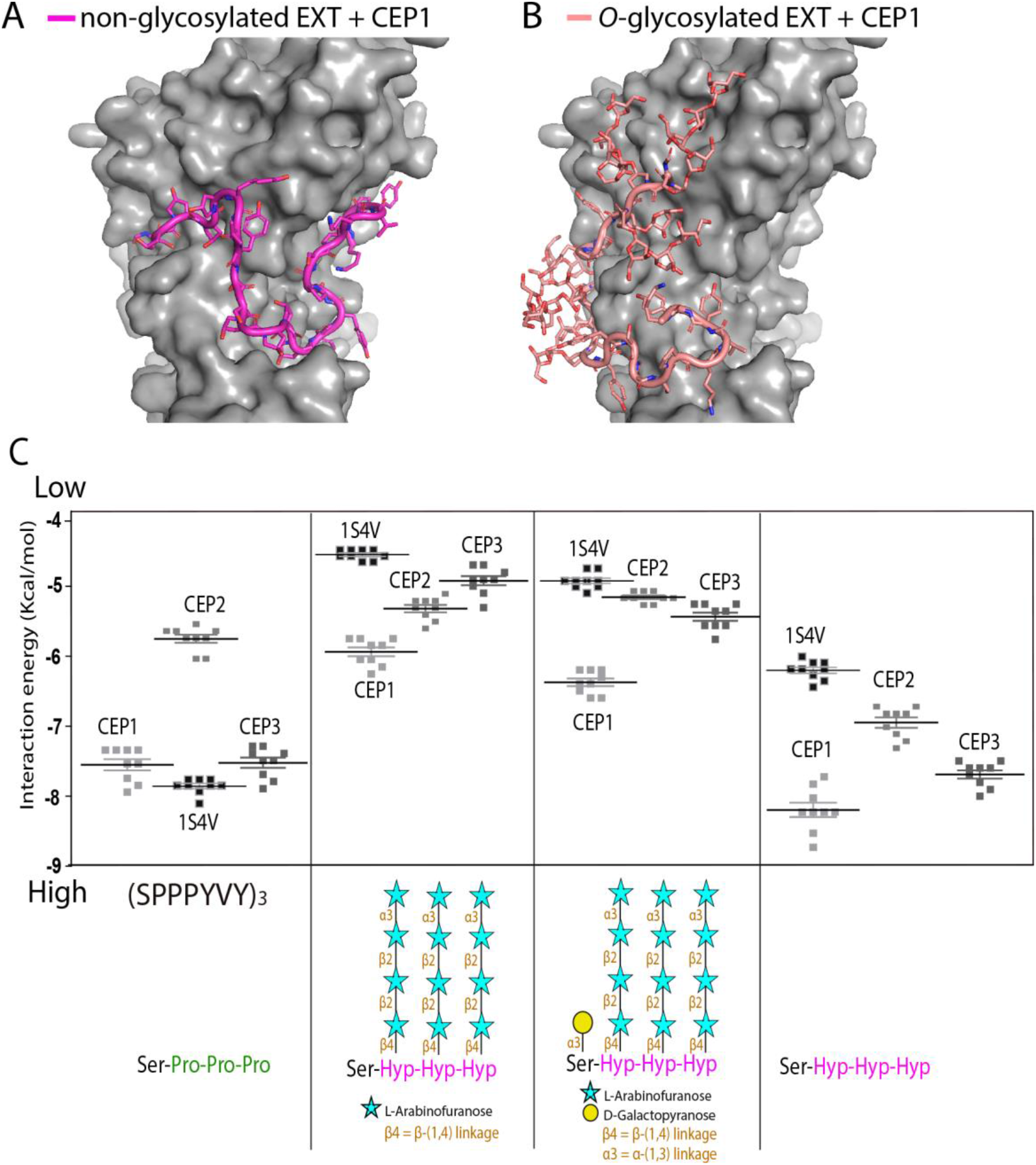
Interaction by an *in silico* docking approach of AtCEP1, AtCEP2 and AtCEP3 with EXT peptides. (**A,B**) Ten docking results for each EXT *O*-glycosylation state are shown superimposed on the AtCEP1 protein surface to evaluate the consistency of docking sites. (**A**) Model of AtCEP1 (protein surface shown in grey) complexed to a non-*O*-glycosylated EXT substrate (SPPPYVY)_3_ (in magenta). (**B**) Model of AtCEP1 (protein surface shown in gray) complexed to an *O*-glycosylated-EXT substrate (protein and *O*-glycans shown in light red, both depicted as sticks). Arabino-galactosylated EXT peptide = [(SOOOYVY)_3_-AG]. (**C**) Comparison of the binding energy of different AtCEP1-AtCEP3 and 1S4V (Ricinus AtCEP included as a control) to EXT substrates with different degrees of *O*-glycosylation. (from left to right) A non-hydroxylated EXT peptide (SPPPYVY)_3_, a hydroxylated but not *O*-glycosylated EXT peptide [(SOOOYVY)_3_; O=hydroxyproline], only arabinosylated EXT-peptide [(SOOOYVY)_3_-A], and arabino-galactosylated EXT peptide [(SOOOYVY)_3_-AG] were analyzed.

**Table S1.**
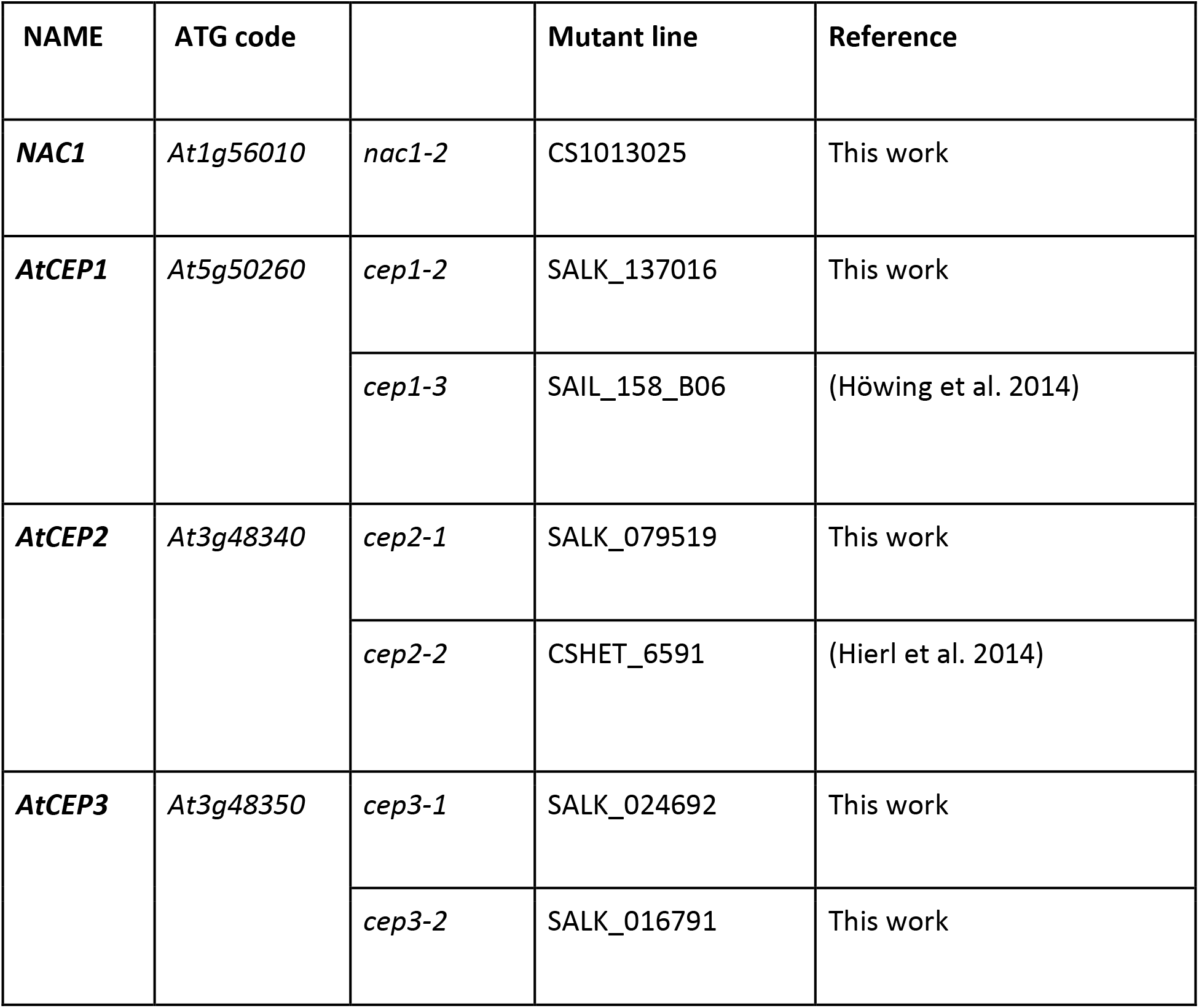
Mutant lines used in this study.

| NAME | ATG code |  | Mutant line | Reference |
| --- | --- | --- | --- | --- |
| ***NAC1*** | *At1g56010* | *nac1-2* | CS1013025 | This work |
| ***AtCEP1*** | *At5g50260* | *cep1-2* | SALK_137016 | This work |
|  |  | *cep1-3* | SAIL_158_B06 | (Höwing et al. 2014) |
| ***AtCEP2*** | *At3g48340* | *cep2-1* | SALK_079519 | This work |
|  |  | *cep2-2* | CSHET_6591 | (Hierl et al. 2014) |
| ***AtCEP3*** | *At3g48350* | *cep3-1* | SALK_024692 | This work |
|  |  | *cep3-2* | SALK_016791 | This work |

**Table S2.** Transgenic lines used in this study.

| NAME | CONSTRUCTION | BACKGROUND | REFERENCE |
| --- | --- | --- | --- |
| **PER8pro::AtCEP2** | promoterPER8::AtCEP2 | Wt Col-0 | (Chen et al. 2016) |
| **PER8pro::AtCEP1** | promoterPER8::AtCEP1 | Wt Col-0 | (Chen et al. 2016) |
| **PER8pro::3×FLAG-NAC1** | promoterPER8::3×FLAG-NAC1 | Wt Col-0 | (Chen et al. 2016) |
| **NAC1pro::NAC1-GUS** | promoterNAC1::NAC1-GUS | Wt Col-0 | (Chen et al. 2016) |
| **AtCEP1pro::3xHA-EGFP-AtCEP1-KDEL** | promoterAtCEP1:: PRE-PRO-3xHA-EGFP-AtCEP1-KDEL | *cep1-3* | (Höwing et al. 2014) |
| **AtCEP1pro::3xHA-EGFP-KDEL** | promoterAtCEP1:: PRE-PRO-3xHA-EGFP-KDEL | *cep1-3* | (Höwing et al. 2014) |
| **AtCEP2pro:: 3xHA-mCherry-KDEL** | promoterAtCEP2::PRE-PRO-3xHA-mCherry-KDEL | *cep2-2* | (Hierl et al. 2014) |
| **AtCEP2pro::3xHA-mCherry -AtCEP2-KDEL** | promoterAtCEP2::PRE-PRO-3xHA-mCherry-AtCEP2KDEL | Wt Col-0 | (Hierl et al. 2014) |
| **AtCEP3pro::GFP** | promoterAtCEP3::GFP | Wt Col-0 | This work |
| **NAC1pro::GFP** | promoterNAC1::GFP | Wt Col-0 | This work |

**Table S3.**
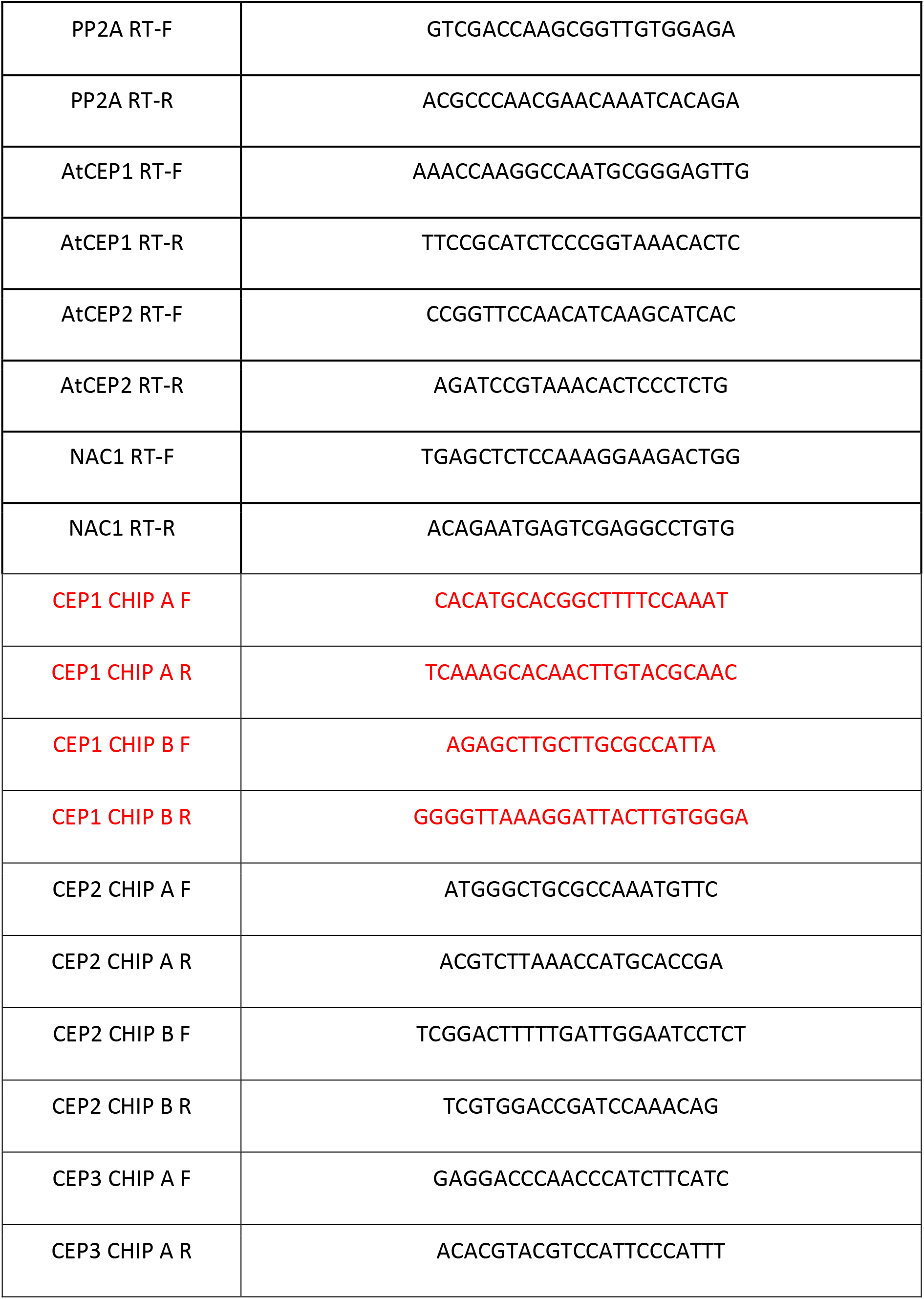
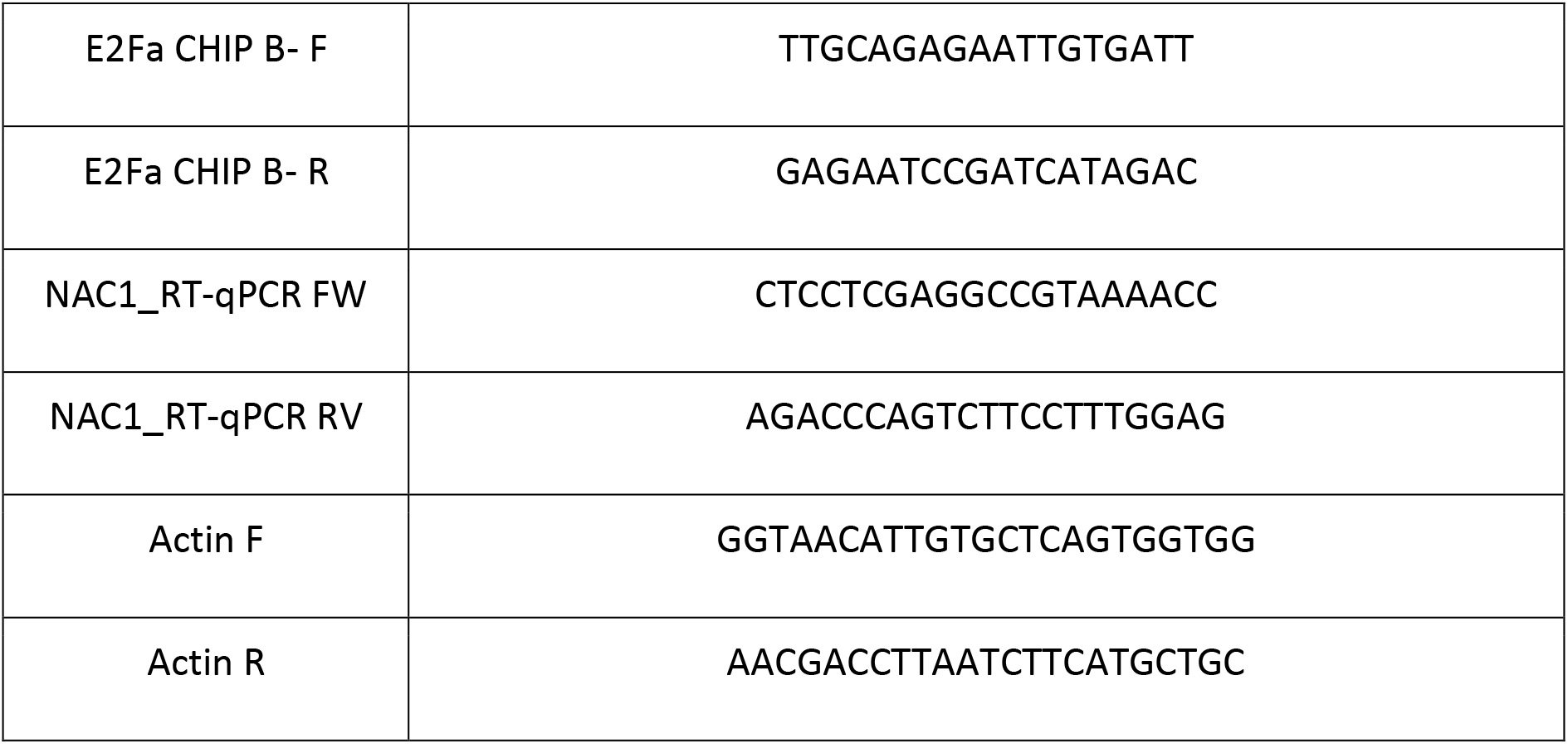
List of primer used.

